# Computational modeling-directed combination treatment with etanercept and mifepristone mitigates neuroinflammation in a mouse model of Gulf War Illness

**DOI:** 10.1101/2025.04.30.651399

**Authors:** Kimberly A. Kelly, Christopher M. Felton, Brenda K. Billig, Ali A. Yilmaz, James P. O’Callaghan, Travis J.A. Craddock, Gordon Broderick, Nancy Klimas, Lindsay T. Michalovicz

**Affiliations:** Health Effects Laboratory Division, National Institute for Occupational Safety and Health, Centers for Disease Control and Prevention, Morgantown, WV, USA; Guest Researcher, Health Effects Laboratory Division, Centers for Disease Control and Prevention, National Institute for Occupational Safety and Health, Morgantown, WV, USA; Department of Biology, Waterloo Institute for Nanotechnology, University of Waterloo, Waterloo, ON, CA; Vaccine and Infectious Disease Organization (VIDO), University of Saskatchewan, Saskatoon, SK, CA; Institute for Neuro-Immune Medicine, Nova Southeastern University, Fort Lauderdale, FL, USA; Department of Clinical Immunology, College of Osteopathic Medicine, Nova Southeastern University, Fort Lauderdale, FL, USA; Miami Veterans Affairs Healthcare System, Miami, FL, USA

**Keywords:** Gulf War Illness, neuroinflammation, therapeutic, anti-inflammatory, anti-glucocorticoid

## Abstract

Gulf War Illness is a chronic multi-symptom disorder experienced by over 30% of veterans from the 1990-1991 Gulf War and is increasingly recognized to be driven by underlying persistent neuroinflammation resulting from chemical and physiological exposures experienced during deployment. Despite significant advances in identifying Gulf War-relevant exposures and underlying pathobiology, effective treatment strategies for Gulf War Illness are still largely lacking. Many studies that have evaluated potential therapies for Gulf War Illness have primarily focused on a single treatment. However, through a mechanistically informed computational evaluation of blood biomarkers and gene expression in veterans with Gulf War Illness, we identified that a combination of anti-inflammatory and anti-glucocorticoid treatment may prove effective in treating Gulf War Illness. Here, we have evaluated combined treatment with the anti-TNFα drug, etanercept, and anti-glucocorticoid, mifepristone, in an established long-term mouse model of Gulf War Illness of combined physiological stress and nerve agent exposure. Supporting results from the computational modeling of this treatment, we found that this drug combination significantly alleviates the underlying neuroinflammation associated with Gulf War Illness. The fusion of computational and *in vivo* preclinical treatment evaluation may provide a highly useful and translationally relevant means by which to identify successful treatment paradigms for Gulf War Illness.

## 1. Introduction

Military warfighters may face many exposures during active duty and deployment that can impact their immediate and long-term health, including burn pits, heat stress, physical encumbrance, excessive exercise, traumatic brain injuries, and chemical exposures. Often the connection between this exposome and veteran health is unclear or poorly understood, making diagnosis and treatment a challenge. Approximately one-third of the veterans of the 1990-91 Gulf War developed a long-term chronic multi-symptom illness, Gulf War Illness (GWI), which has been connected with several potential in-theater exposures – most notably, sarin and cyclosarin nerve agents and other organophosphate compounds (1–3). A significant amount of work, both preclinical and clinical/epidemiological, has been invested in understanding the underlying mechanisms of GWI with the major goal of identifying better treatments to control or eliminate symptoms. Numerous studies have correlated the neuroinflammatory consequences of these exposures and long-term neuroinflammation and aberrant inflammatory processes in ill veterans (4–10). In particular, several studies have shown that combined exposure to high physiological stress and Gulf War-relevant organophosphates or nerve agent surrogates is associated with long-term neuroinflammatory sensitization (11–16). As these potential underlying disease mechanisms have been identified, subsequent investigations have focused on treatments targeting neuroinflammatory processes in the brain to treat GWI. As such, several studies have identified the potential for botanicals, nutraceuticals, and existing pharmaceuticals to reduce neuroinflammation in various animal models of GWI (15,17–28).

Previous work in animal models has demonstrated that exposure to high levels of physiological stress mimicked by chronic exogenous exposure to corticosterone stress hormone exacerbates the neuroinflammatory response to the organophosphate sarin surrogate, diisopropyl fluorophosphate (DFP) (11–14). Moreover, this initial exposure primes the brain for hyperreactivity to future inflammatory challenges, particularly when kindled by intermittent re-exposure to stressor episodes (15,16). These observations in the animal model align with computational models of GWI that describe an aberrant, homeostatic neuroimmune state in veterans with GWI (4,29,30). By employing these neuroimmune interaction models in treatment analysis, several potential therapeutic interventions have been identified (29–32). Due to the complexity of the proposed pathological interactions associated with GWI, many of these strategies have identified combination treatments targeting different molecular mechanisms associated with GWI to achieve the highest level of symptom remission. While many previous treatment strategies for GWI have specifically focused on managing specific sets of symptoms, these combination treatments focus specifically on targeting the underlying disease mechanisms of GWI. Specifically, many of the treatment strategies targeted neuroendocrine and immune dysregulation to achieve the highest levels of remission, identifying a need to target cytokine and glucocorticoid signaling (29–32).

Based on the outcomes of this numerical modeling and the corresponding computer simulated treatment courses, we hypothesized that a combination therapy targeting Th1 cytokines and the glucocorticoid receptor employed after the induction of GWI would reduce the severity of the exacerbated neuroinflammatory response to subsequent systemic inflammatory challenge in our established long-term GWI mouse model (15). In this study, mice were treated with etanercept, a tumor necrosis factor (TNF) antagonist, and mifepristone, a progesterone antagonist, one to two weeks following the initial CORT DFP exposure either during or outside of the subsequent alternating CORT exposure in the 5-week protocol, similar to previous work (15). Treatment efficacy was determined as the drug combination’s ability to reduce the exacerbated neuroinflammatory response to subsequent LPS challenge across multiple brain areas. Treatment with etanercept and mifepristone during CORT exposure significantly reduced the expression of cytokine mRNA in the cortex, hippocampus, and striatum compared to CORT DFP LPS exposed mice that did not receive treatment. These results support the hypothesis that treatment with etanercept and mifepristone resets the aberrant homeostatic neuroinflammatory signaling associated with GWI and observed in our long-term GWI models (15,16), normalizing the response to future inflammatory challenges.

## 2. Materials and Methods

### 2.1 Materials

The following drugs and chemicals were provided by or obtained from the sources indicated: DFP, LPS and mifepristone (Sigma-Aldrich, St. Louis, MO, USA), Etanercept (Enbrel, Amgen, Thousand Oaks, CA, USA), and CORT (Steraloids, Inc., Newport, RI, USA). All other reagents and materials were of at least analytical grade and were obtained from a variety of commercial sources.

### 2.2 Animals and long-term GWI exposure paradigm

All animal procedures were performed within protocols approved by the US Centers for Disease Control and Prevention-Morgantown Institutional Animal Care and Use Committee (13-JO-M-021) and the US Army Medical Research and Development Command Animal Care and Use Review Office (GW120045.01) in an AAALAC International accredited facility. Upon arrival, adult male C57BL/6J mice (N=26; 6-8 weeks old, ∼30g; The Jackson Laboratory, Bar Harbor, ME, USA; RRID:IMSR_JAX:000664) were single housed and allowed to acclimate for at least one week prior to the exposures. Mice were given food (Harlan 7913 irradiated NIH-31 modified 6% rodent chow) and water ad libitum and received daily health checks from animal husbandry personnel.

The mice were exposed to our established long-term GWI mouse model (15). Briefly, mice received CORT (200mg/L in 0.6% EtOH) in the drinking water for 7 days followed by a single injection of DFP (4 mg/kg, i.p, ∼ LD_25_) on Day 8. Due to the potential toxicity of DFP, animals were observed and monitored following DFP exposure and evaluated by activity scoring once at approximately 6 hrs post-exposure and twice daily for 2 additional days. Activity scoring was defined as follows: 4, no voluntary movement when cage is opened; 3, no voluntary movement when gently stroked; 2, delayed slow escape behavior when gently stroked; 1, slow voluntary escape behavior without touching; 0, normal movement and escape behavior on cage lid opening. Any animal that presented with a non-zero activity score or displaying other adverse effects of exposure (e.g., intensely labored breathing, complete immobility) by 48 hrs post-DFP, conditions indicative of a higher risk of mortality, were euthanized by CO_2_ inhalation or pentobarbital-based euthanasia solution (100-300 mg/kg, i.p., Fatal Plus, Vortech Pharmaceuticals, Dearborn, MI, USA).

Following this initiating exposure, mice were re-exposed to CORT water every other week for 4 weeks and then challenged with LPS (0.50 mg/kg, s.c.) on Day 35. Mice were sacrificed by decapitation 6 hours post-LPS or saline (0.9%, controls) exposure. Prior to dosing, mice were randomly assigned to one of 4 treatment groups: Saline (N=5), GWI (CORT DFP LPS, N=7), GWI + Treatment during CORT (N=7), and GWI + Treatment outside CORT (N=7). The N/group was increased for all GWI groups to account for potential mortality due to cholinergic toxicity (∼25%).

### 2.3 Etanercept/mifepristone treatment

Previous computational modeling studies predicted that combined, tiered treatment with a T-helper type 1 (Th1) cytokine inhibitor and subsequently with a glucocorticoid receptor (GR) inhibitor could recover normal neuroendocrine-immune function in a high proportion of veterans with GWI (4,31,32). Following advanced discussions and additional modeling (see 33–35), etanercept and mifepristone were selected for the Th1 and GR inhibitors, respectively. The timing of administration for the combination treatment was based upon the tiered treatment strategy described previously (32) and the half-life of the drugs in mice: 4 days for etanercept and 5 hours for mifepristone; etanercept was given 2 days prior to mifepristone.

Mice received the etanercept (5 mg/kg, i.p.) and mifepristone (20 mg/kg, i.p.) combination treatment during the second week of CORT (Day 16 and 18 for etanercept and mifepristone, respectively) (Figure 1A) or the following week without CORT (Day 23 and 25) (Figure 1B). Doses and route of exposure for etanercept and mifepristone were chosen following literature review and subsequent dose response studies (data not shown). Briefly, mice were given etanercept (2.5-20 mg/kg, i.p.) 24 hours prior to LPS (2.0 mg/kg, s.c.) and inflammatory cytokine mRNA responses were measured at 2 and 6 hours post-LPS to assess anti-inflammatory responses of etanercept. For mifepristone, mice received CORT for 7 days, as previously described, with mifepristone (10-40 mg/kg, i.p.) given on Day 4 followed by LPS on Day 7; inflammatory cytokine mRNA responses were measured at 6 hours post-LPS to assess the ability of mifepristone, an anti-GR drug, to reduce the glucocorticoid-primed inflammatory response to LPS.

**Figure 1.**
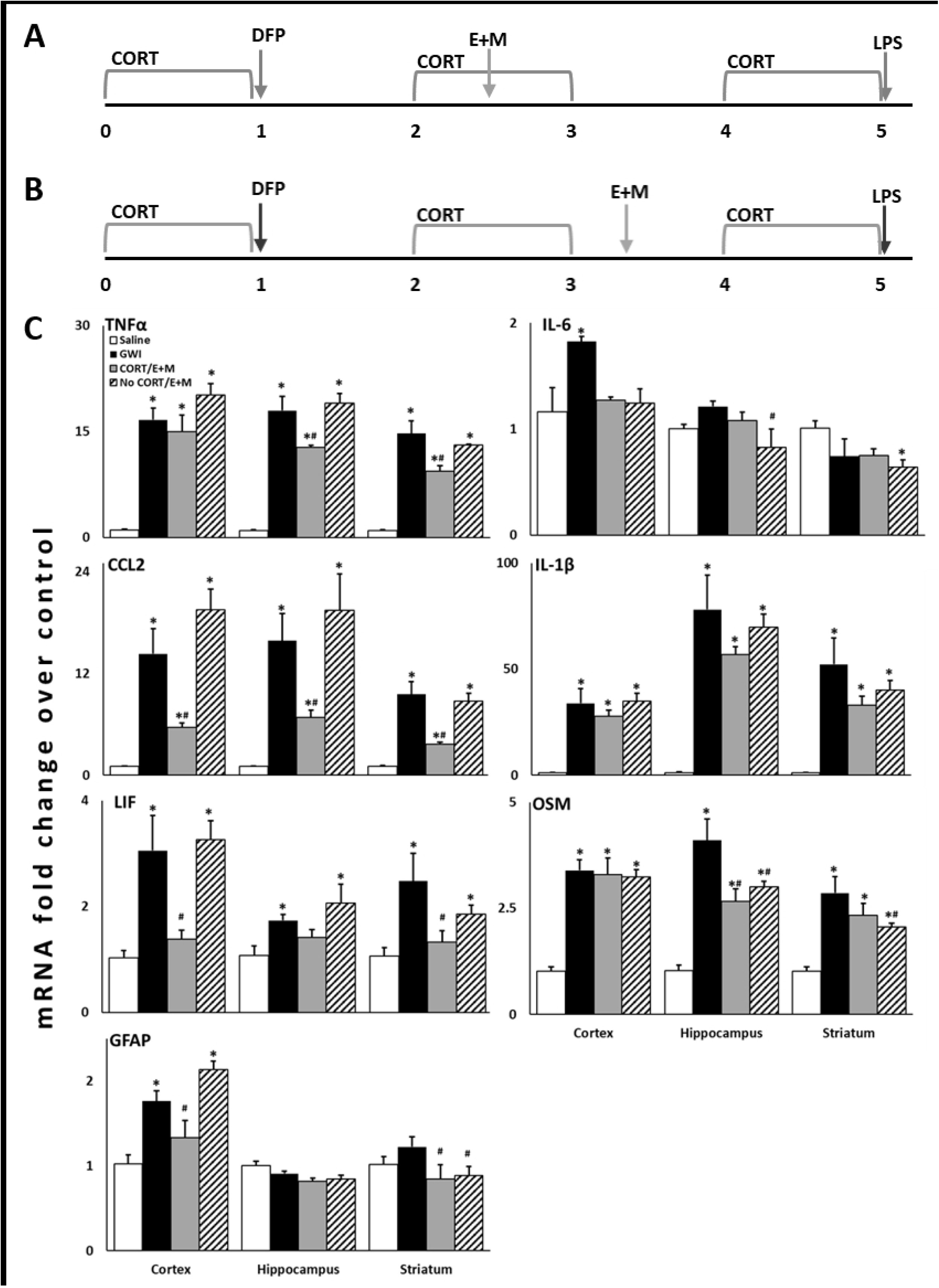
Combination treatment with etanercept and mifepristone reduces the expression of inflammatory cytokines in the brain resulting from Gulf War Illness exposure-related neuroinflammation. Mice (*N*=4-5 mice/group) were given corticosterone (CORT; 200 mg/L in 0.6% ethanol) in the drinking water for seven days followed by a single injection of diisopropyl fluorophosphate (DFP; 4 mg/kg, i.p.) on Day 8. These mice were then given CORT every other week for an additional 4 weeks with an LPS challenge (0.5 mg/kg, s.c.) on Day 35. Mice receiving combined treatment with etanercept and mifepristone (E+M) received the drugs spaced 2 days apart either during the 2^nd^ week of CORT (A: etanercept, Day 16; mifepristone, Day 18; CORT/E+M) or between the 2^nd^ and 3^rd^ weeks of CORT (B: etanercept, Day 23; mifepristone, Day 25; No CORT/E+M). Brain areas were collected at 6 hours post-LPS exposure (Day 35) for qPCR analysis. C) The cortex, hippocampus, and striatum were evaluated for changes in the mRNA expression of several inflammatory cytokines. *P* ≤ 0.05 vs. *Saline or ^#^CORT DFP LPS (GWI). Bars represent mean ± S.E.M.

### 2.4 Brain dissection and tissue preparation

Immediately after decapitation, whole brains were removed from the skull with the aid of blunt curved forceps. Cortex, hippocampus, and striatum were dissected free hand on a thermoelectric cold plate (Model TCP-2, Aldrich Chemical Co., Milwaukee, WI, USA) using a pair of fine curved forceps (Roboz, Washington, DC, USA). Brain regions were frozen at −85 °C and used for subsequent isolation of total RNA.

### 2.5 Real-time qPCR

Total RNA from cortex, hippocampus, and striatum was isolated as previously described (12,36). Real-time PCR analysis of the housekeeping gene, glyceraldehyde-3-phosphate dehydrogenase (*Gapdh*), and of the proinflammatory mediators, tumor necrosis factor-α (*Tnf*), interleukin-6 (*Il6*), C-C motif chemokine ligand 2 (*Ccl2*), interleukin-1β (*Il1b*), leukemia inhibitor factor (*Lif*), and oncostatin M (*Osm*) was performed in an ABI7500 Real-Time PCR System (Thermo Fisher Scientific, Waltham, MA, USA) in combination with TaqMan® chemistry, as previously described (12,36). Relative quantification of gene expression was performed using the comparative threshold (ΔΔCT) method. Changes in mRNA expression levels were calculated after normalization to GAPDH. The ratios obtained after normalization are expressed as fold change over corresponding saline-treated controls.

### 2.6 Statistical analysis

Sample size was determined to be N=4/group based on previous calculations (12). A larger N/group was used to account for DFP-related mortality and statistical outliers. Statistical outliers were identified using Grubb’s test, as previously described (15,37). Statistical significance was determined per cytokine between groups within each brain area by one-way ANOVA and overall treatment effects across all brain areas and cytokines by three-way ANOVA (Cytokine x Exposure x Brain Area) of log-transformed values using SigmaPlot v15 (Systat Software, Inc., San Jose, CA, USA; RRID:SCR_003210) followed by Fisher LSD post-hoc analysis (P ≤ 0.05). All data are available in the NIOSH Data and Statistics Gateway (https://www.cdc.gov/niosh/data).

## 3. Results

A combination treatment of etanercept and mifepristone was given at two different time points in our long-term GWI mouse model (15): one during CORT exposure (Figure 1A) and one outside of CORT exposure (Figure 1B). Previously, we have shown that this long-term model represents a dormant inflammatory phenotype instigated by the initial CORT+DFP exposure that results in an exacerbated neuroinflammatory response to subsequent systemic inflammatory challenge with LPS (15,16). Here, we found that the etanercept/mifepristone combination treatment significantly reduced the expression of several cytokines across the three different brain areas that were evaluated (Figure 1C). Treatment during CORT significantly reduced inflammatory cytokine mRNA expression across all brain areas compared to the GWI exposure (*P* < 0.001), whereas treatment outside of the CORT exposure only had a significant effect in the striatum (*P* = 0.015). Specifically, etanercept/mifepristone treatment significantly reduced *Tnf* expression in the hippocampus and striatum, *Ccl2* expression in all three brain areas, *Lif* expression in the cortex and striatum, and *Gfap* expression in the cortex when administered during CORT. In addition, *Osm* expression in the hippocampus and *Gfap* expression in the striatum were significantly reduced by the treatment regardless of whether it was given during or outside of CORT exposure. Striatal *Osm* was the only mRNA that was exclusively reduced by treatment given outside of the CORT exposure.

## 4. Discussion

Through a significant research effort over the last several years, it has been shown that GWI is associated with underlying neuro-immune dysfunction and several constituents of the Gulf War theatre exposome, particularly high physiological stress and organophosphate nerve agent or pesticides, have been demonstrated to impact neuroinflammation in animal models. Specifically, our CORT DFP model has been shown to produce significant exacerbation of neuroinflammatory responses, both acutely and long-term, as well as altered brain structure and behavioral responses relevant to structural changes and cognitive impairments measured in veterans with GWI (11,12,14–16,26). Cutting-edge computational modeling, rooted in mechanistic prior knowledge and first principles of regulatory biology, has allowed for the translational validation of this GWI model, as well as provided predictions for complex, multi-drug strategies to treat GWI (30–35). These studies largely indicated a tiered and time-lagged, multi-drug approach targeting aberrant immune responses and endocrine disruption identified in veterans with GWI as a therapeutic approach with a high probability for achieving symptom remission (32). Here, we utilized a combination treatment of the Th1 cytokine antagonist, TNF inhibitor, etanercept, with the glucocorticoid inhibitor, mifepristone. By evaluating how this treatment impacted the previously established exacerbation of inflammatory cytokine mRNA expression in the brain in our long-term GWI model, we demonstrated that etanercept/mifepristone combination therapy was able to reduce the neuroinflammatory profile across multiple areas of the brain.

Over the last decade, several animal models have been developed for GWI to capture different aspects of the exposome. Some of the most well-studied models include chronic exposure to low-level organophosphate, combined with exposure to the nerve agent prophylactic, pyridostigmine bromide, and pyrethroid insecticide, permethrin, with or without stressor exposure. The corticosterone and organophosphate exposure models used in this study (6,7), also have subsequently been used to evaluate potential treatments. These models have evaluated varying cellular, molecular, and behavioral outcomes of exposure, including neuroinflammation.

Accordingly, several recent studies have investigated the potential for treatments to improve underlying neuroinflammation in these models, particularly in reducing brain cytokine expression and activation of microglia and astrocytes – hallmarks of neuroinflammation (15,17–28). However, the approach to evaluating these treatments has varied across the studies, which can complicate evaluating their translational potential. While some studies applied treatments simultaneous to the GW-relevant model exposures, curcumin, melatonin, lacto-N-fucopentaose III (LNFPIII), propranolol, and the etanercept-mifepristone combination assessed in the current study were provided after the initiating exposures in the respective GWI animal models (15,19,20,26).

While the former studies may provide insights into preventing the development of exposure-related disease, the latter better replicate the conditions of ill veterans who have already been exposed and are over 30 years removed from the initiation of illness. This highlights the importance of considering the timing of treatment administration in animal models of human illness to increase the potential successful translation of these therapies into ill veterans.

The long-term CORT DFP GWI rodent model was developed to encapsulate both the in-theatre exposure conditions of high physiological stress and neurotoxic organophosphate exposure experienced by veterans during deployment, as well as the intermittent stressors and symptom flare-ups that would be experienced by ill veterans following their deployment (15,16). Therefore, this model combines two different, but crucial, aspects of GWI: 1) the purported initiating exposures of illness (CORT DFP), and 2) the illness-related inflammatory symptom presentation (modeled by LPS challenge). As described previously, this model represents an underlying or dormant primed neuroinflammatory condition (15). Specifically, animals exposed to CORT DFP and subsequent waves of CORT exposure demonstrate baseline, normal levels of cytokine mRNA in the brain unless challenged with an inflammatory stimulant, such as LPS. When an inflammatory challenge is given, however, the resulting response is significantly exacerbated compared to both controls and CORT LPS exposed animals. In developing a treatment paradigm within the long-term GWI mouse model, the goal was to provide treatment at a time point that would not interfere with the instigation of GWI, i.e. removed from the initial CORT DFP exposure, as well as not directly treat the inflammatory challenge. It is this lagged effect and the temporal evolution of response to subsequent triggers that are uniquely capture by this mechanistic analysis of homeostatic regulatory processes. In the absence of a specific treatment for GWI, therapeutic management largely relies on addressing and reducing specific symptoms. Thus, the goal of the computational studies directed at treatment identification for GWI focused on achieving stable remission through regulatory reprogramming of the underlying aberrant disease processes (32). In the animal model, this was translated to preventing the exacerbated response to inflammatory challenge through immune-modulation, as opposed to abolishing all normal inflammatory processes or immunosuppression. Therefore, the combination treatment was given 10-17 days prior to the LPS challenge (see Figure 1A and B). The observation that treatment with etanercept and mifepristone significantly reduced cytokine mRNA expression across various brain regions compared to the untreated GWI group while still producing an inflammatory response distinguishable from saline controls suggests that treatment may have targeted or “reset” the underlying aberrant neuroinflammatory phenotype.

The development of treatments for CNS illnesses and diseases is extremely challenging and many therapies fail during development (38,39). Often drugs targeting neurological diseases have promising preclinical results but fail to demonstrate efficacy in clinical trials potentially due to the availability of relevant animal models, clear targets for treatment, among other factors (38). Here, along with previously published work, a treatment discovery pipeline was developed and utilized that shows promise in addressing several of the hurdles involved in treating neuro-based illness and disease. This pipeline utilized a variation of a bedside-to-bench-to-bedside approach to identify a potential effective combination treatment for GWI. This was achieved through a mechanistically driven computational modeling of human disease aligned to explain patient-collected biomarker data (4,31) and the evaluation of treatments through large-scale computer simulation of strategies directed at delivering a “reset” of aberrant processes (31,32). This was followed by preclinical evaluation of treatment potential in an established/translational animal model (Figure 1) and ultimately leading to entrance of the treatment strategy into clinical trial (Pilot: NCT04255498; Follow-up: NCT04254627) (Figure 2). This knowledge-driven treatment design framework addresses several of the factors identified in the review from Gribkoff and Kaczmarek (38) that can be applied to the preclinical aspect of treatment development, such as the use of a translationally-relevant preclinical model of GWI (30), a multi-system hypothesis related to disease mechanisms of GWI (i.e. endocrine and neuro-immune), as well as the application of a combination therapy. The collective work of the current and previous studies indicates the value of combining computational and *in vivo* clinical evaluation of potential drugs to strengthen the preclinical support of therapeutic strategies.

**Figure 2.**
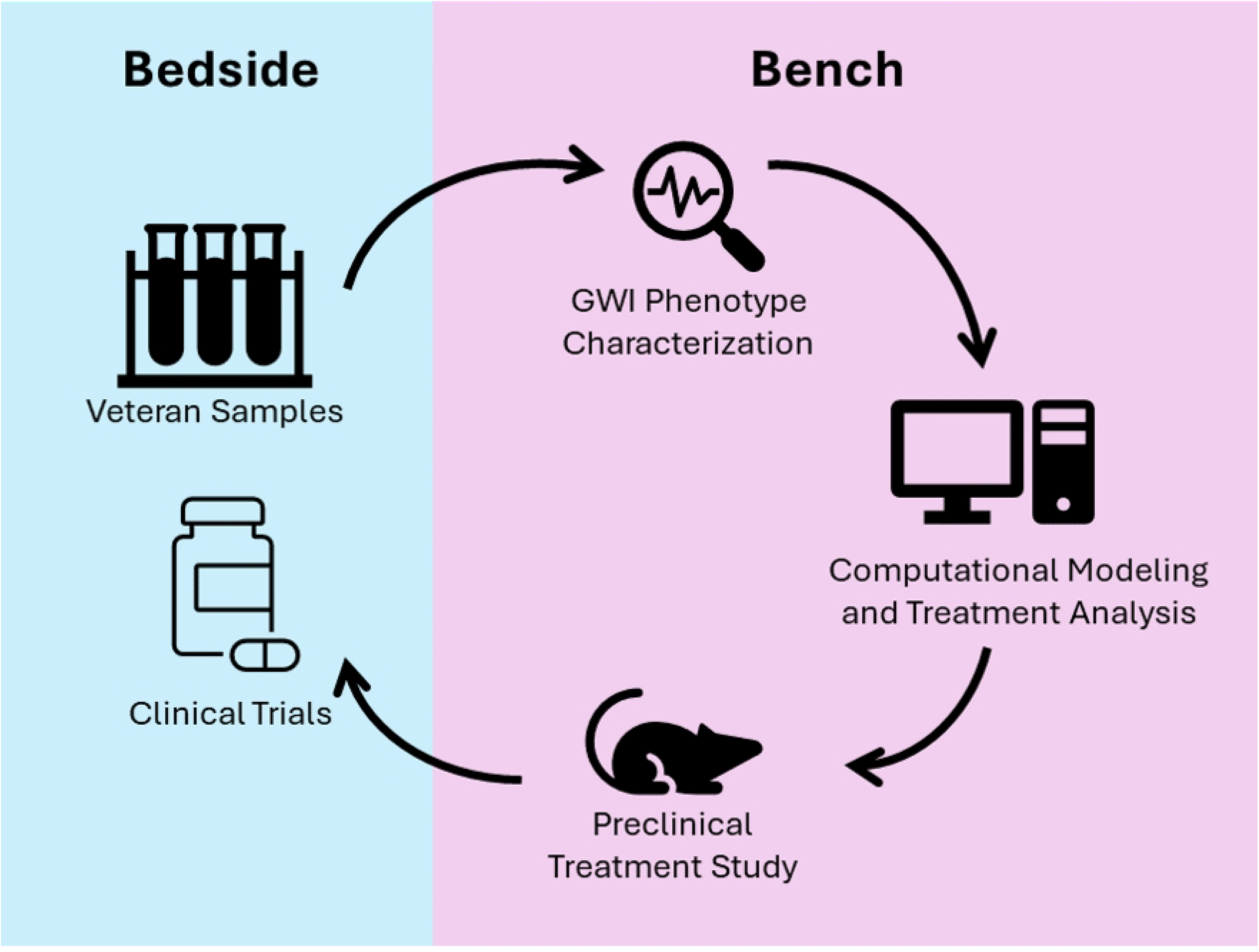
Bedside-to-Bench-to-Bedside Pipeline for Treatment Identification and Evaluation. Through a collaborative consortium effort, veteran biosamples were collected and subjected to multi-modal analysis, i.e. RNAseq and blood chemistry. These data were subsequently utilized to build a logic model of the key molecules and signaling mechanisms represented in the samples from veterans with GWI. This logic model was then used to evaluate and develop potential treatment strategies, including combination therapy, to repurpose FDA-approved drugs to reset the GWI phenotype described in the model. The resultant hypothesized combination treatment strategy was then developed for and tested in an existing animal model of GWI to evaluate its efficacy in reducing GWI-associated neuroinflammation. The combined results of *in silico* and *in vivo* preclinical testing of the treatment strategy supported subsequent clinical trial development and execution.

While the etanercept and mifepristone combination provide promising preclinical results for the potential efficacy of this treatment strategy in addressing the underlying pathobiology of GWI, there are a few limitations to the current study. It is important to note that the preclinical treatment evaluation was performed using only male animals. While most veterans with GWI are male, studies have indicated differences in GWI profiles between males and female veterans (40–43). While the acute CORT DFP mouse model of GWI has been evaluated in female mice (44), the long-term GWI mouse model has not been characterized in female mice. Future work would address the evaluation and validation of the long-term GWI mouse model in female mice against female veterans with GWI, along with preclinical evaluation of the efficacy of the etanercept and mifepristone treatment with careful consideration of the implications of utilizing a progesterone blocker in females. In addition, the current study evaluates the therapeutic effects of the combination treatment solely on the basis of cytokine mRNA expression changes. While previous work has correlated the cytokine mRNA changes in the acute and long-term GWI models to brain cellular and structural changes, downstream neuroinflammatory signaling pathways, and peripheral inflammatory markers (11–14,16), additional follow-up with behavioral and peripheral measures in animals would add to a greater understanding of the extended impact of the treatment on multiple factors related to GWI presentation. Overall, this study presents *in vivo* preclinical data in support of a computational-derived combination therapy for GWI and highlights a pathway for the identification and evaluation of potential treatments for GWI.

## Acknowledgements

This work was supported by a CDMRP GWIRP research award [W81XWH-13-2-0085; Klimas, Morris, PIs] and intramural funds from the CDC, National Institute for Occupational Safety and Health (NIOSH; O’Callaghan, Kelly, Michalovicz, PIs). VIDO receives operational funding from the Canada Foundation for Innovation through the Major Science Initiatives Fund and from the Government of Saskatchewan through Innovation Saskatchewan and the Ministry of Agriculture (Broderick). This research was also supported, in part, thanks to funding from the Canada Research Chairs Program [CRC-2022-00204; Craddock PI] and the University of Waterloo (Craddock).

The findings and conclusions in this report are those of the authors and do not necessarily represent the official position of the National Institute for Occupational Safety and Health, Centers for Disease Control and Prevention. This work was supported by the Assistant Secretary of Defense for Health Affairs through the Gulf War Illness Research Program. Opinions, interpretations, conclusions, and recommendations are those of the author and are not necessarily endorsed by the Department of Defense. This article is published with the permission of the Director of VIDO, journal series no. TBD.

## Author Contributions

K.K., J.O., T.C., G.B., N.K., and L.M. conceptualized the project. K.K., J.O., T.C., N.K., and L.M. were responsible for funding acquisition. K.K., J.O., and L.M. were responsible for methodology, project administration, supervision, validation, and resources for the presented study. K.K., C.F., B.B., A.Y., and L.M. were responsible for data curation and investigation. L.M. was responsible for visualization of the data and preparing the original manuscript draft. All authors reviewed and edited the manuscript.

## Notes

### Competing Interest Statement

The authors have declared no competing interest.

